# Metformin strongly affects gut microbiome composition in high-fat diet-induced type 2 diabetes mouse model of both sexes

**DOI:** 10.1101/2020.10.30.362095

**Authors:** Laila Silamiķele, Ivars Silamiķelis, Monta Ustinova, Zane Kalniņa, Ilze Elbere, Ramona Petrovska, Ineta Kalniņa, Jānis Kloviņš

## Abstract

Effects of metformin, the first-line drug for type 2 diabetes therapy, on gut microbiome composition in type 2 diabetes have been described in various studies both in human subjects and animals. However, the details of the molecular mechanisms of metformin action have not been fully understood. Moreover, there is a significant lack of information on how metformin affects gut microbiome composition in female mice models, as most of the existing studies have focused on males only.

Our study aimed to examine metformin-induced alterations in gut microbiome diversity and composition of high-fat diet-induced type 2 diabetes mouse model, employing a randomized block, factorial study design, and including 24 experimental units allocated to 8 treatment groups. We performed shotgun metagenomic sequencing using DNA obtained from fecal samples representing gut microbiome before and after ten weeks-long metformin treatment.

We identified 100 metformin-related differentially abundant species in high-fat diet-fed mice before and after the treatment, with most of the species abundances increased. In contrast, no significant changes were observed in control diet-fed mice.We also observed sex-specific differences in response to metformin treatment. Males experienced more pronounced changes in metabolic markers, while, in females, the extent of changes in gut microbiome representatives was more marked, indicated by 53 differentially abundant species with more remarkable Log fold changes compared to the combined-sex analysis. Our results suggest that both sexes of animals should be included in future studies focusing on metformin effects on the gut microbiome.

## Introduction

Metformin is the first-line therapy for the treatment of type 2 diabetes (T2D). According to the American Diabetes Association and European Association for the study of Diabetes guidelines, it is the preferred option for initiating glucose-lowering due to its efficacy, safety, tolerability, and low cost (Davies et al., 2018). The molecular mechanism of action of metformin, however, remains unclear.

It is generally considered that metformin’s antihyperglycemic is mainly due to reduced hepatic glucose production, thus activating pathways independent of AMP-activated protein kinases (AMPKs) (Wu et al., 2017). There is increasing evidence that metformin’s action mechanism is associated with the physiological processes in the gastrointestinal tract. For example, a more pronounced effect of metformin can be observed when the drug is administered orally than intravenously at an equivalent dose (Stepensky et al., 2002). It has been estimated that 20-30% of people receiving metformin therapy develop gastrointestinal side effects, with approximately 5% being unable to tolerate metformin at all (Dujic et al., 2016). Metformin accumulates in gastrointestinal tissues (Kinaan et al., 2015); for example, it is 30-300 times more concentrated in the small intestine than in plasma, and 30-50% of the drug reaches the colon and is eliminated with feces (Graham et al., 2011).

Studies in humans have convincingly shown that metformin specifically alters gut microbiome (Forslund et al., 2015; Wu et al., 2017). It has been shown that metformin exerts its effects on the microbiome already in the first 24 hours (Elbere et al., 2018) and that the alterations in the gut microbiome could be related to the metformin-induced immune response (Ustinova et al., 2020).

A number of studies that have specifically targeted metformin effects in rodents have used 16S rRNA sequencing and have identified that metformin modifies the metabolic profile of HFD-fed animals accompanied with changes in the microbiome (Ji et al., 2019; H. Lee et al., 2018; Zhang et al., 2015).

We have employed an experimental design that includes both sexes’ mice, comparing metformin effects in HFD, and adequately composed control diet-fed mice. We also use shotgun metagenome sequencing providing better taxonomical resolution, thus, for the first time, presenting valuable information on sex-related differences in response to metformin treatment regarding changes in gut microbiome composition.

## Materials and Methods

### Study design

We designed this study as a randomized block experiment comprised of three blocks with a three-way factorial treatment arrangement where factors of interest are T2D status induced by high-fat diet (HFD) or control diet (CD) feeding, sex, and metformin therapy status, forming eight different treatment groups – HFD_M_Met−, HFD_F_Met−, HFD_M_Met+, HFD_F_Met+, CD_M_Met−, CD_F_Met−, CD_M_Met+, and CD_F_Met+ (Figure 1). In each of the eight treatment groups, we included three experimental units, totaling 24. Study’s sample size was determined by the Resource Equation method, appropriate for complex designs. As the experimental unit was a cage with three animals – the total number of animals involved in the experiment was 72.

**Figure 1.**
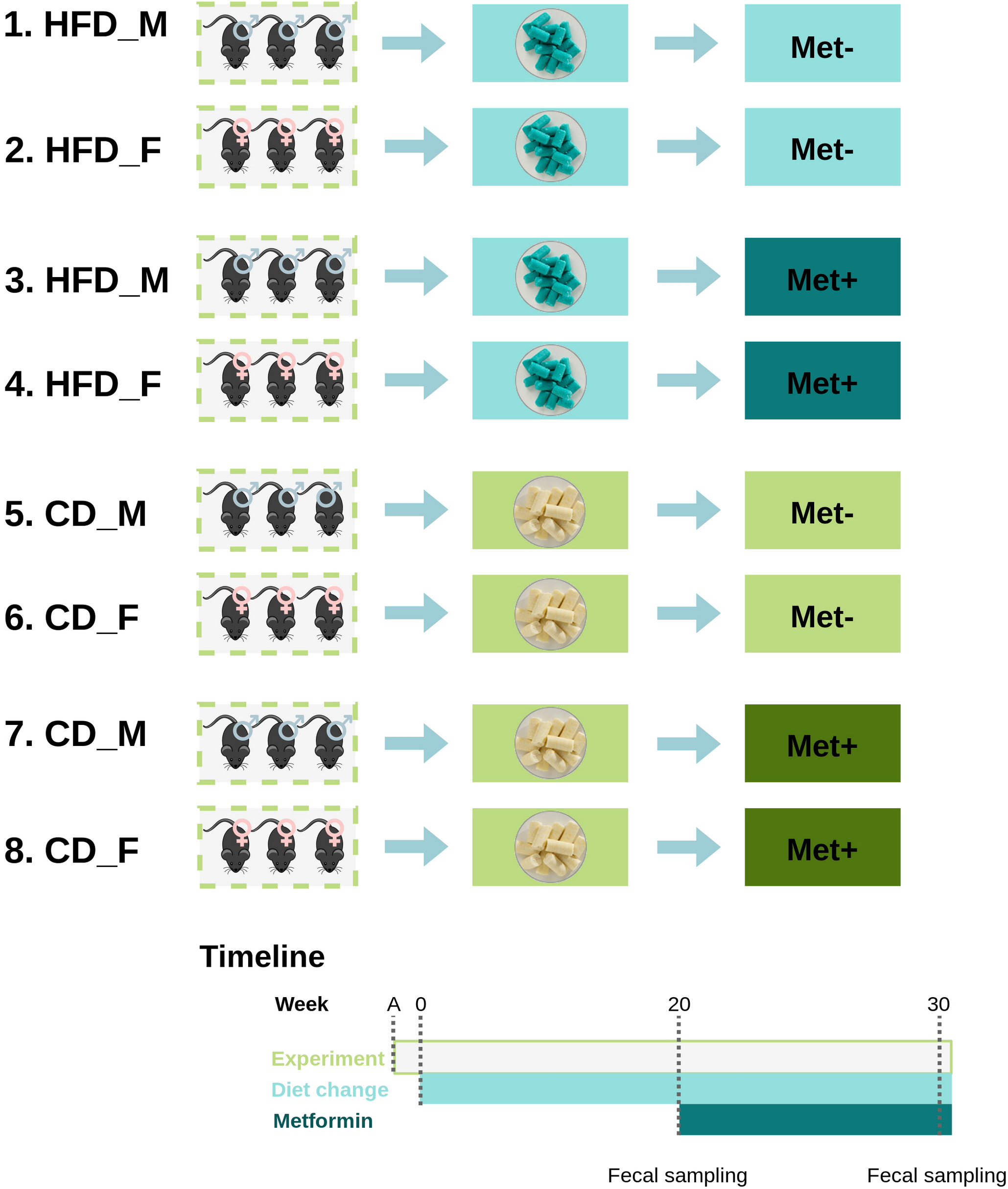
Experimental design and timeline of the study. Experiment was performed in 3 randomized blocks each separated in time. Each block contained 1 experimental unit (cage with 3 animals of the same sex) representing one of the 8 experimental groups: 1) HFD_M_Met−; 2) HFD_F_Met−; 3) HFD_M_Met+; 4) HFD_F_Met+; 5) CD_M_Met−; 6) CD_F_Met−; 7) CD_M_Met+; 8) CD_F_Met+. Abbreviations for the factors: HFD – high-fat diet-fed mice; CD – control diet-fed mice; M – male mice; F – female mice; Met− – mice not receiving metformin treatment; Met+ – mice receiving metformin treatment. After 1 week long adaptation period experimental units were randomized into HFD- or CD-fed groups. After 20 weeks an onset of T2D manifestations was observed, first collection of fecal samples performed and each of the groups was randomized into Met− and Met+ groups for the duration of 10 weeks which was followed by the endpoint of the study and second collection of fecal samples.

After the adaptation week, all experimental units of the same block were randomly assigned to HFD- or CD-fed groups so that each of the treatment groups would consist of animals with similar body weight, and experimental units of both sexes would be represented in the same number in both types of treatment groups. After the induction of T2D manifestations, experimental units were randomly assigned to receive or not to receive metformin treatment providing that the number of experimental units in each of the groups is equal. During all the procedures, treatments, measurements, and sample collection were performed randomly within the same block. Work with each of the blocks was performed on separate days of the week, keeping the interval between interventions in the same block constant. Blinding was applied where appropriate.

### Experimental animals

Age-matched 4-5-week-old male and female C57BL/6N mice corresponding to specific pathogen-free (SPF) status were obtained from the University of Tartu Laboratory Animal Centre.

All the animals were housed in SPF conditions, 23 ± 2 °C, 55% humidity. The light cycle was 12:12 hours, with a light period from 7:00 am to 7:00 pm. All the procedures were performed during the first half of the day in a specially designated procedure room.

Animals were housed in individually ventilated cages (Techniplast) up to three same-sex animals per cage on aspen bedding mixed with ALPHA-dri. All the animals had free access to drinking water. Animals were fed HFD or CD *ad libitum*.

All the cages were enriched with carton tunnels, plastic sheds, wooden sticks, and nesting material. For the whole duration of the experiment, animals were observed once a day; if we observed any type of severe suffering that could not be alleviated, the suffering animal was euthanized by cervical dislocation.

### Experimental procedures

After a one-week long adaptation period during which animals were fed regular chow diet *ad libitum* and received regular drinking water a diet change was initiated. All the cages from each of the blocks were randomly assigned to a high-fat diet-fed group or control diet-fed group. Animals were provided with a rodent diet containing 60 kcal% fat (D12492, Research Diets) or rodent diet with 10 kcal% fat (D12450J, Research Diets) *ad libitum*. Both types of diets were sterilized by irradiation. Body weight and food intake (per cage) were measured every week; water intake (per cage) – twice a week. Four weeks after starting the assigned diet, we initiated a measurement of blood glucose levels at two weeks interval using an Accu-Chek Performa glucometer (Roche) after fasting for 6 hours. To estimate the insulin resistance, HOMA-IR index was calculated (by formula 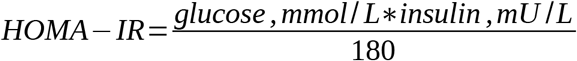) based on fasting plasma glucose and insulin levels determined by the mouse glucose assay and mouse insulin ELISA kit (both from Crystal Chem) at week 20.

The cages were changed every week. Upon opening the cage, each mouse was immediately transferred to a clean, separate box in which animals were allowed to defecate voluntarily. Each of the animals was weighted. Feces were collected in sterile tubes in three aliquots. Bottles with drinking water were changed two times a week.

Blood for regular glucose level measurements was obtained from the saphenous vein by puncturing the vein with 25G needle. Accu Chek Performa glucometer with Accu Chek test strips was used to measure the glucose level in blood. Before the procedure animals were fasted for 6 hours starting from 8 am to 14 pm. The induction of type 2 diabetes was evaluated by glucose and insulin level measurements in plasma. The plasma necessary for the analysis was obtained from blood drawn from the saphenous vein. The maximum volume drawn at a one-time point was in line with generally accepted guidelines.

Metformin was provided to mice with drinking water. The concentration of metformin was calculated to correspond to 50 mg/kg body mass/day. During the therapy period, all of the bottles, including those of the control group, were changed every day. Metformin was freshly added to the drinking water every day upon water changing. The duration of metformin therapy was ten weeks.

All the animals were sacrificed by cervical dislocation without any other anesthesia, as the effect of other medications would interfere with the study’s aims.

### Microbial DNA isolation and shotgun metagenomic sequencing

The DNA from fecal samples representing each experimental unit at two timepoints – before and after the metformin treatment (N = 48) was extracted with the FastDNA®SPIN Kit for Soil. DNA yield was determined using the Qubit dsDNA HS Assay Kit on the Qubit® Fluorometer 1.0 (Invitrogen Co.). The extracted DNA samples were then diluted to a concentration of 5 ng/μl.

Libraries for metagenomic shotgun sequencing were prepared using MGIEasy Universal DNA Library Prep Kit (MGI Tech Co., Ltd.); the construction of libraries was completed in a single batch. The input of DNA was 200 ng. Library preparation was performed according to the Universal DNA Library Prep Set User Manual and spiked with 1% PhiX. For metagenomic analysis each experimental unit was represented by one randomly chosen animal as the microbiome is shared between animals in the same cage.

Pooled, circularized and barcoded libraries were used as templates for DNA nanoball preparation (16 samples per lane) and further analyzed on DNBSEQ-G400RS next generation sequencing platform (MGI Tech Co., Ltd.) using DNBSEQ-G400RS High-throughput Sequencing Set (FCL PE150) (MGI Tech Co., Ltd.) according to manufacturer’s instructions. Sequencing depth was calculated to achieve at least 20 million paired-end 150 bp reads per sample.

### Data analysis

Read quality evaluation was performed with FastQC (Andrews, 2010). Adapter clipping was performed with cutadapt v1.16 (Martin, 2011). Reads were trimmed from 5’ and 3’ end using 5 bp window with quality threshold 20 using Trimmomatic v0.39 (Bolger et al., 2014). Paired reads with length 75 bp or longer were retained for further data processing.

Reads originating from the host were removed by mapping reads against mouse reference genome GRCm38 release 96. Taxonomic classification of unmapped reads was performed with Kraken 2.0.8-beta against progenomes database (Mende et al., 2020) with additionally included mouse (GRCm38) and human (GRCh38) reference genomes (Wood et al., 2019). Only reads with a confidence score of 0.5 or higher were regarded as classified. Abundance reestimation was done with bracken v2.5 at species level (Lu et al., 2017). Reads classified as *Homo sapiens* or *Mus musculus* were removed from subsequent analyses. Taxonomies with low read counts were removed using filterByExpr function implemented in edgeR 3.26.8 (Robinson et al., 2009).

Due to the complex experimental design, the differential abundance analysis was not performed using statistical tools that take into account compositional nature of the data. Differential abundance testing was performed with limma 3.40.6 using voom transformation with sample quality weights (Ritchie et al., 2015). Differential testing was performed for combinations of multiple factors: metformin usage, time, diet, and sex. Correction for multiple testing was implemented with Benjamini-Hochberg method. Taxa with FDR ≤ 0.05 were regarded statistically significant. The effect of individual mice was accounted using duplicateCorrelation function in limma.

Alpha diversity was expressed as the exponential of Shannon diversity index resulting in the effective number of species, genera, and phyla at the respective taxonomic levels.

To account for the compositional nature of taxonomic data, the imputation of zero values was performed with Bayesian-multiplicative replacement method as implemented in R 3.6.3 package zCompositions 1.3.4 with default parameters (Martín-Fernández et al., 2015). Resulting taxonomic data were subsequently transformed using centered log ratio transformation with scikit-bio 0.5.5 (Aitchison, 1986).

Aitchison’s distance was used as beta diversity metric. Principal component analysis biplot was constructed from the transformed compositions with scikit-learn 0.22.

Differences in body weights and biochemical parameters between HFD-fed and CD-fed groups were determined by one-way anova and t-test. The normality of measurement distributions was assesed by Shapiro-Wilk test, equality of variances was evaluated by an F-test.

Hypothesis testing for changes in alpha diversity before and after therapy was performed using paired t-test for each metformin and diet group separately. The normality of the diversity differences between time points was assessed with the Shapiro-Wilk test. P-values < 0.05 were considered statistically significant.

## Results

### The effect of a high-fat diet on body weight and metabolic parameters of mice

Mice were fed with HFD for 20 weeks in order to induce T2D manifestations. Significant differences in body weights between HFD-fed and CD-fed mice were observed after two weeks with mean body weight 22.89 ± 3.36 and 19.9 ± 3.08 g, respectively (P-value = 0.03) (Figure 2). As expected, body weight was higher in males than in females in each of the diet groups. Metformin treatment had no significant effect on body weight gain in any of the studied groups.

**Figure 2.**
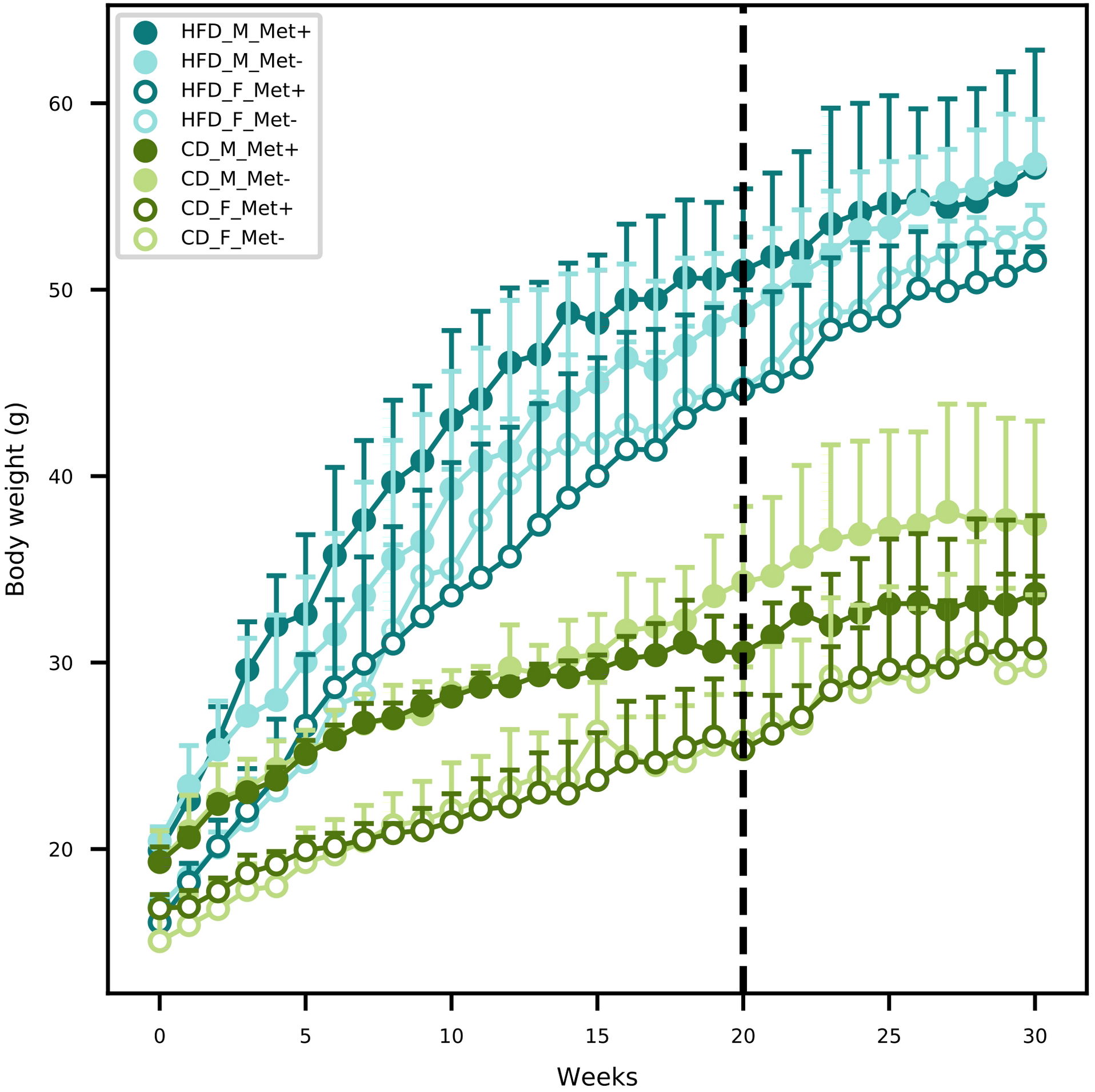
Mean body weight of each of the experimental groups in each of the weeks. Dashed line indicates the beginning of metformin treatment.

Fasting blood glucose level was monitored by glucometer fortnightly. Upon detecting statistically significant differences in blood glucose levels between HFD-fed and CD-fed groups, we initiated regular determination of fasting glucose and insulin levels in plasma samples (Table 1). We calculated HOMA-IR index, and values above two were considered to correspond to insulin resistance suggesting the onset of T2D. Before the beginning of the metformin treatment mean HOMA-IR index was above 2 in all of the HFD-fed groups but below 2 in all of the CD-fed groups. After the ten-week-long metformin treatment, fasting glucose and insulin levels in plasma samples were remeasured.

**Table 1.**
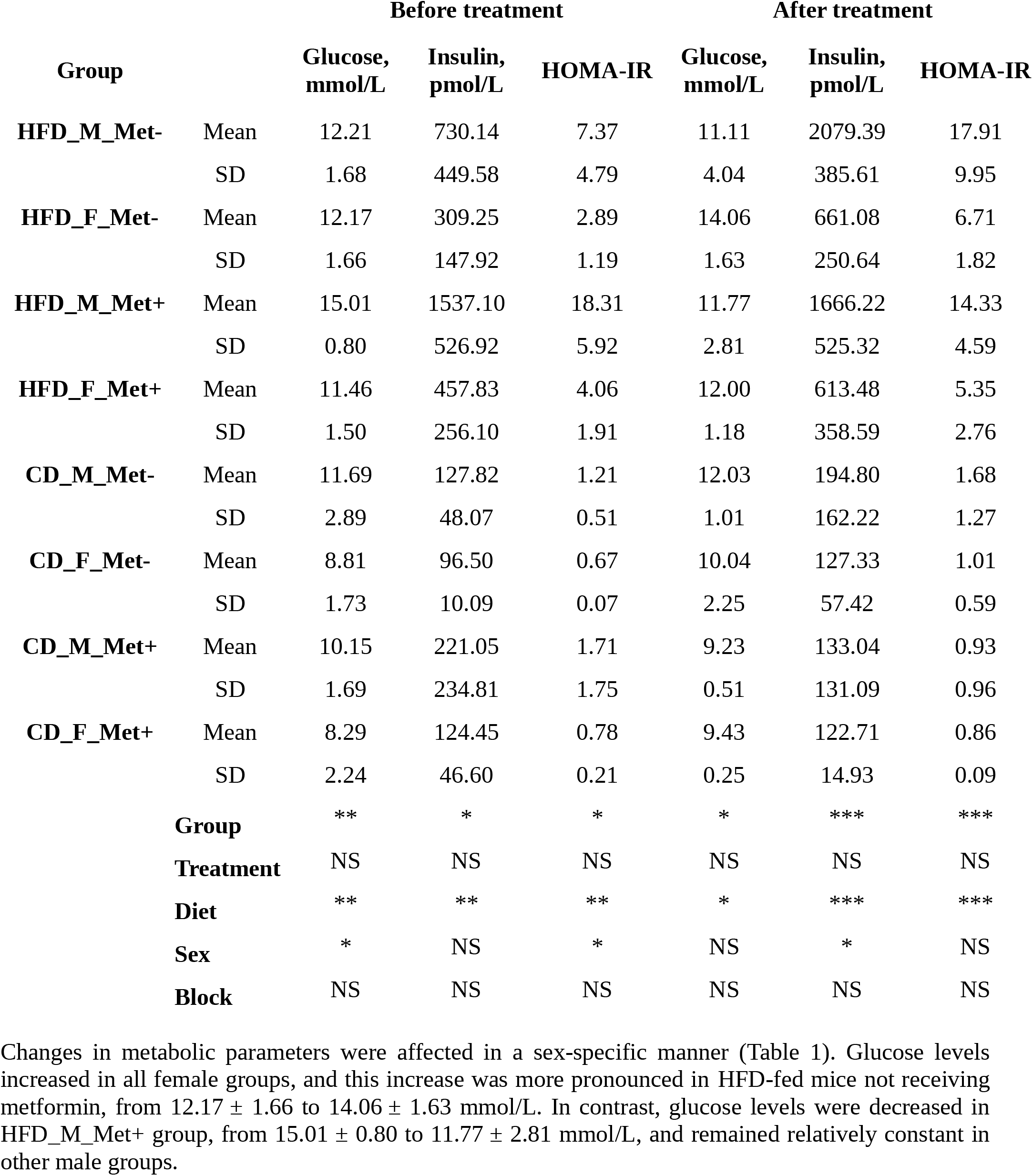
Biochemical parameters before and after metformin treatment. N = 3 in each of the studied groups. Significance codes: 0 (***), 0.001 (**), 0.01 (*), >0.05 (NS).

|  |  | Before treatment |  |  | After treatment |  |  |
| --- | --- | --- | --- | --- | --- | --- | --- |
| Group |  | Glucose,<br>mmol/L | Insulin,<br>pmol/L | HOMA-IR | Glucose,<br>mmol/L | Insulin,<br>pmol/L | HOMA-IR |
| **HFD_M_Met-** | Mean | 12.21 | 730.14 | 7.37 | 11.11 | 2079.39 | 17.91 |
|  | SD | 1.68 | 449.58 | 4.79 | 4.04 | 385.61 | 9.95 |
| **HFD_F_Met-** | Mean | 12.17 | 309.25 | 2.89 | 14.06 | 661.08 | 6.71 |
|  | SD | 1.66 | 147.92 | 1.19 | 1.63 | 250.64 | 1.82 |
| **HFD_M_Met+** | Mean | 15.01 | 1537.10 | 18.31 | 11.77 | 1666.22 | 14.33 |
|  | SD | 0.80 | 526.92 | 5.92 | 2.81 | 525.32 | 4.59 |
| **HFD_F_Met+** | Mean | 11.46 | 457.83 | 4.06 | 12.00 | 613.48 | 5.35 |
|  | SD | 1.50 | 256.10 | 1.91 | 1.18 | 358.59 | 2.76 |
| **CD_M_Met-** | Mean | 11.69 | 127.82 | 1.21 | 12.03 | 194.80 | 1.68 |
|  | SD | 2.89 | 48.07 | 0.51 | 1.01 | 162.22 | 1.27 |
| **CD_F_Met-** | Mean | 8.81 | 96.50 | 0.67 | 10.04 | 127.33 | 1.01 |
|  | SD | 1.73 | 10.09 | 0.07 | 2.25 | 57.42 | 0.59 |
| **CD_M_Met+** | Mean | 10.15 | 221.05 | 1.71 | 9.23 | 133.04 | 0.93 |
|  | SD | 1.69 | 234.81 | 1.75 | 0.51 | 131.09 | 0.96 |
| **CD_F_Met+** | Mean | 8.29 | 124.45 | 0.78 | 9.43 | 122.71 | 0.86 |
|  | SD | 2.24 | 46.60 | 0.21 | 0.25 | 14.93 | 0.09 |
| **Group** |  | ** | * | * | * | *** | *** |
| **Treatment** |  | NS | NS | NS | NS | NS | NS |
| **Diet** |  | ** | ** | ** | * | *** | *** |
| **Sex** |  | * | NS | * | NS | * | NS |
| **Block** |  | NS | NS | NS | NS | NS | NS |

Insulin level increased in all female groups (except CD_F_Met+ where it remained constant), similar to what was observed with glucose level. In males, insulin level elevated in all but CD_M_Met+ group. In HFD_M_Met− group, the increase was statistically significant, from 730.14 ± 449.58 to 2079.39 ± 385.61 pmol/L (P-value < 0.05), whereas the increase in HFD_M_Met+ group was minor.

We detected changes in the HOMA-IR index in all HFD-fed groups (Table 1). In HFD_M_Met− and HFD_F_Met− groups, insulin resistance increased over time in females reaching statistical significance (P-value < 0.05). Unexpectedly, however, in HFD-fed groups receiving metformin, the effect was different for males and females. Metformin decreased insulin resistance in males, from 18.31 ± 5.92 to 14.33 ± 4.59, but in females, the treatment had no diminishing effect – on the contrary, the HOMA-IR index was increased in the HFD_F_Met+ group, from 4.06 ± 1.91 to 5.35 ± 2.76, though not statistically significant. In CD-fed groups, we observed a similar effect of metformin treatment – in CD_M_Met− and CD_F_Met− groups and CD_F_Met+ group HOMA-IR index grew over time, but in males receiving the treatment, it declined.

### Differences in microbiome composition between experimental groups

We determined the microbiome composition of fecal samples by shotgun metagenomic sequencing. The median value of the obtained paired-end reads was 40044941 (IQR 10957336). After quality trimming and host removal, a median of 28807601 (IQR 9664060) reads were retained. The median percentage of classified reads after taxonomic classification was 40.33% (IQR 12.24%).

The abundances of the top genera in each of the experimental groups are depicted in Figure 3. For comprehensibility reasons, only the genera with a relative abundance of at least 1% are presented. Before the initiation of the treatment, in all HFD-fed groups, *Lactococcus, Bacteroides*, and genera representing *Lachnospiraceae* dominated the gut microbiome composition. Other top taxa represented in all HFD-groups were *Muribaculaceae*, *Lactobacillus*, *Parabacteroides*, *Mucispirillum*, and *Dorea*. *Bacteroides* prevailed in all CD-fed groups, followed by *Lactococcus, Lachnospiraceae*, and *Muribaculaceae*, the abundance of which was greater in all CD-fed groups compared to HFD-fed mice.

**Figure 3.**
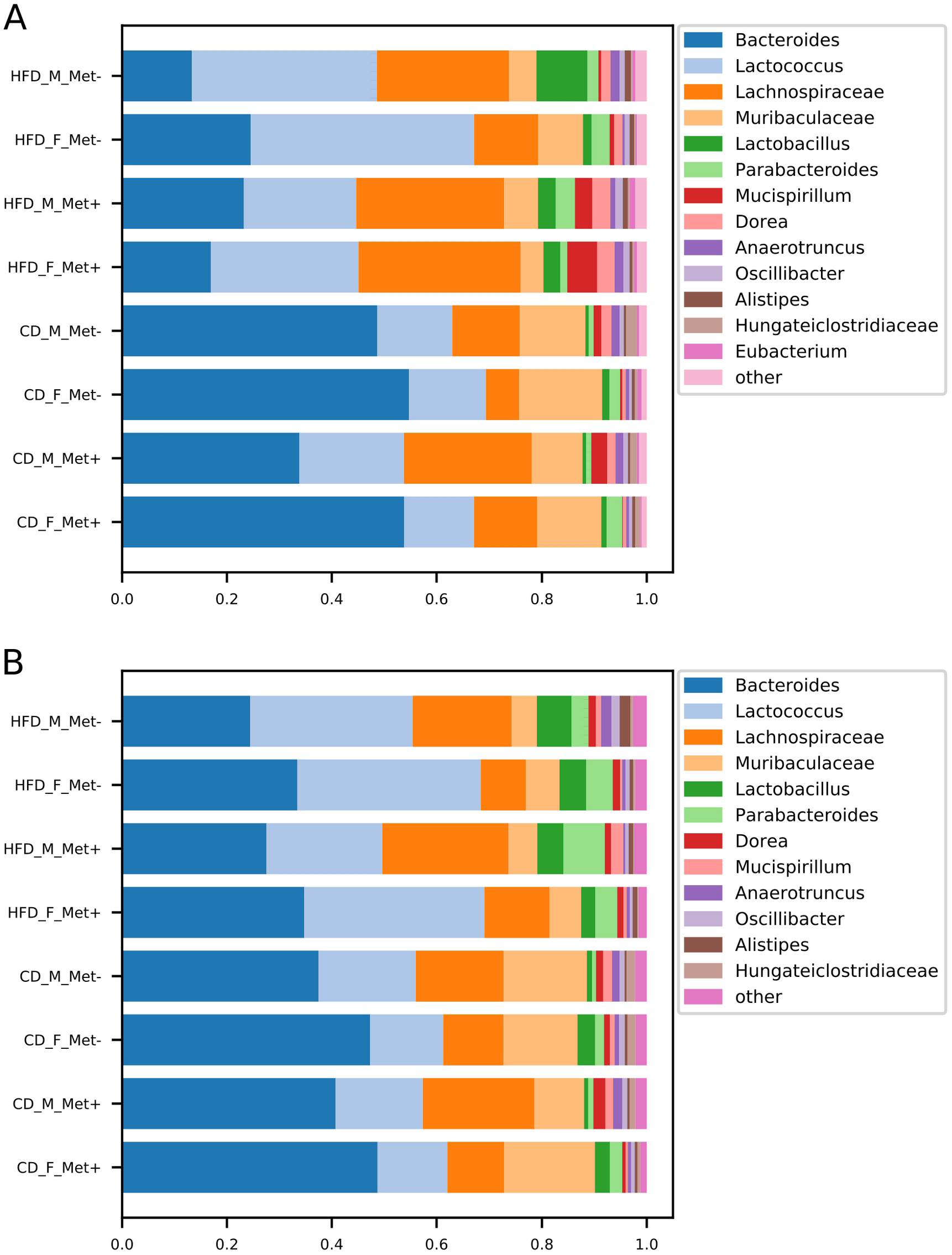
Microbiome composition at genus level in each of the experimental groups before and after the treatment. The abundances of top genera are expressed as proportions, only the genera or other lowest identified taxa with the relative proportion of at least 1% are shown. (**A**) time point before the treatment; (**B**) time point after the treatment.

At the time point after the treatment, we identified an increase in *Bacteroides* abundance in all HFD-fed groups, which was maintained at the expense of reduced abundance of *Lachnospiraceae*, whereas *Bacteroides*, *Lactococcus*, and *Lachnospiraceae* remained top taxa in all HFD-fed groups. All CD-fed groups were dominated by *Bacteroides*, followed by smaller proportions of *Lactococcus*, *Lachnospiraceae*, and *Muribaculaceae*.

### Diversity analysis

#### Alpha diversity

We identified a decrease in the alpha diversity of the HFD_Met+ groups before and after metformin treatment (Figure 4), though not statistically significant. In CD_Met+ groups and groups that did not receive metformin, no alpha diversity changes were observed before and after metformin treatment. Overall the effective number of species (ENS) was higher in HFD-fed groups – mean ENS before treatment was 10.99 ± 3.04 and 9.62 ± 2.64 in HFD-fed and CD-fed mice, respectively. The same was observed at the time point after the treatment – mean ENS was 10.62 ± 3.55 and 10.18 ± 1.70 in HFD-fed and CD-fed mice, respectively.

**Figure 4.**
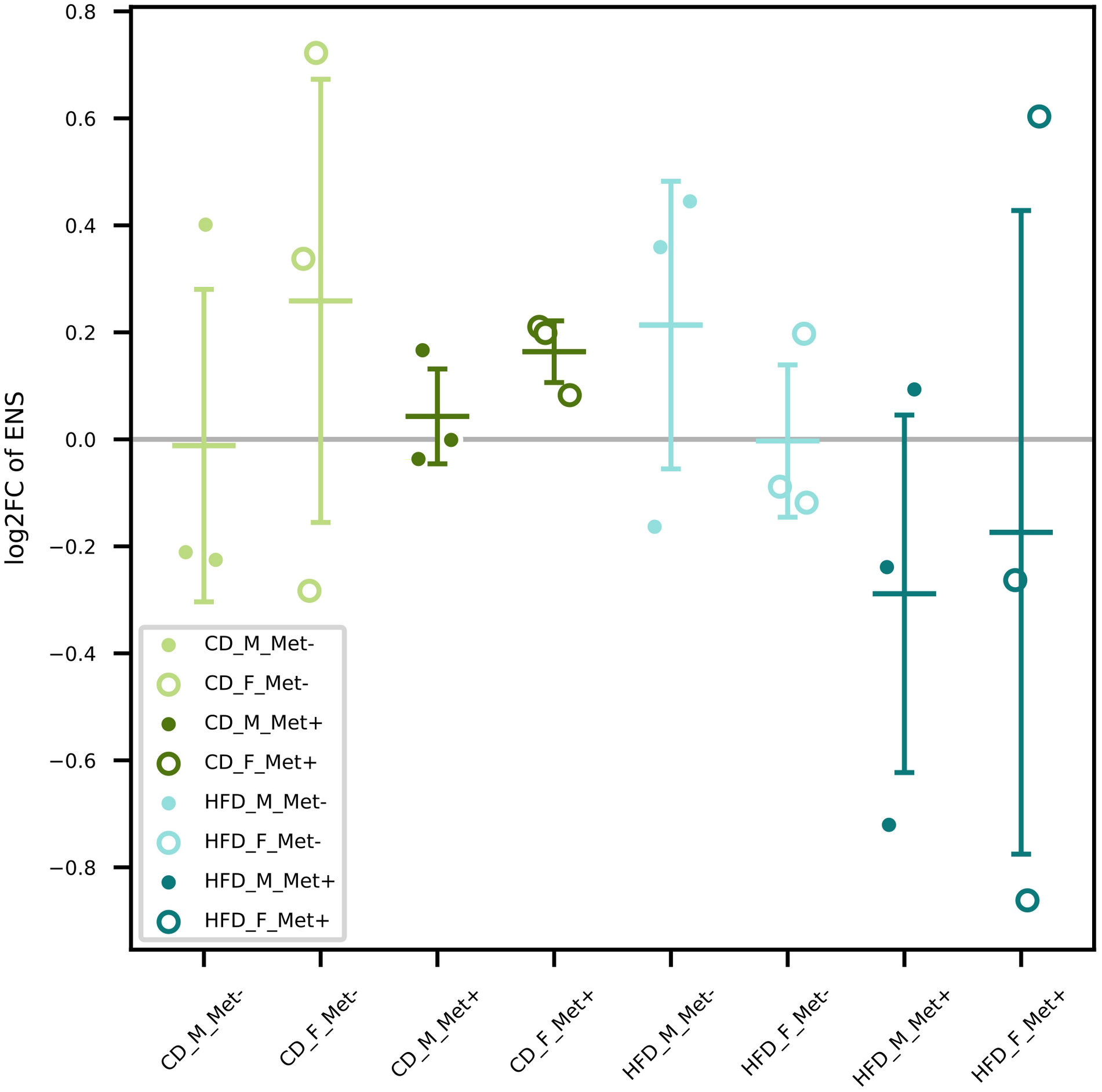
Changes in alpha diversity in each of the groups expressed as effective number of species.

#### Beta diversity

Beta diversity analysis revealed clustering depending on various factors included in the study (Figure 5). Samples obtained from HFD-fed mice clustered apart from those of CD-fed mice at both time points, with the principal microbial identifiers being *Pediococcus acidilactici*, *Lactobacillus spp.*, *Desulfomicrobium orale*, *Desulfovibrio fairfieldensis*, *Fusicatenibacter saccharivorans*, *Proteus spp.* before the beginning of metformin treatment. *Lactospiraceae bacterium 6_1_37FAA*, *Staphylococcus xylosus*, *Intestinimonas massiliensis* supplemented this list of microbial identifiers after the metformin treatment, though some species were represented by < 100 reads in any of the samples. Furthermore, when analyzing both sexes separately, before the treatment, HFD_F groups were directed toward *Lactobacillus* vectors, but HFD_M groups towards *Proteobacteria* members. In both sexes of CD-fed mice, principal identifiers were affiliated to *Bacteroidetes* and *Clostridia*, although the species were different for males and females. After the treatment, samples of HFD-fed mice tended to cluster closer, and *Clostridia* members appeared among the most characteristic taxa of these groups. In CD_F groups, representatives of *Bacteroidetes* and *Clostridia* remained the principal identifiers; however, in CD_M_Met+ group, a shift towards *Proteobacteria* was observed. Nevertheless, no apparent clustering regarding metformin treatment status was observed.

**Figure 5.**
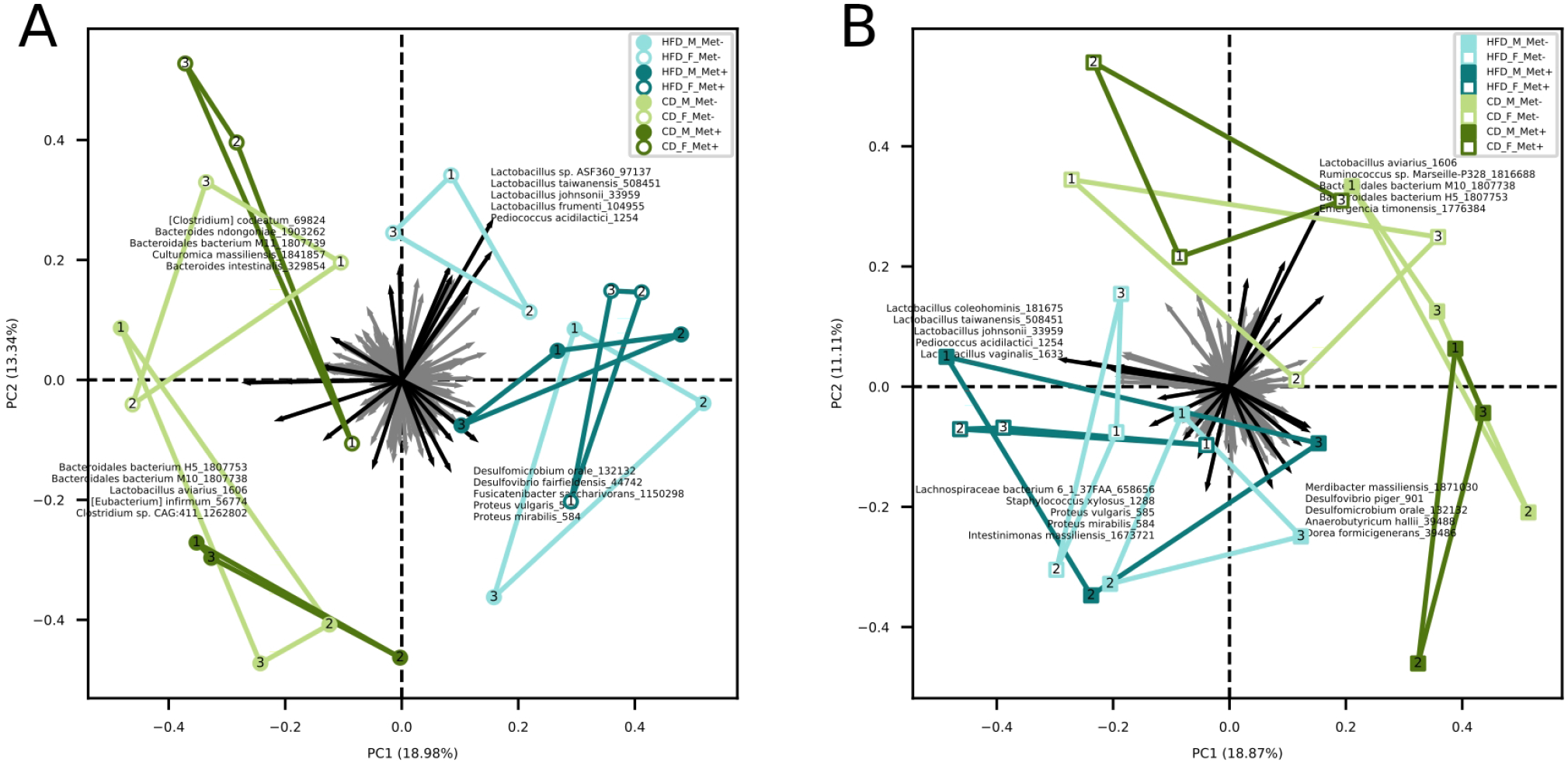
Biplots indicating the beta diversity in each of the groups. (**A**) Beta diversity before the beginning of metformin treatment. (**B**) Beta diversity after 10 weeks long metformin treatment. Samples representing each experimental unit in each of the experimental groups are shown as dots.

### Differentially abundant species between treatment arms

We evaluated the abundance of different species of microbes between groups using various contrasts shown in Figure 6 (see supplementary data for further details).

**Figure 6.**
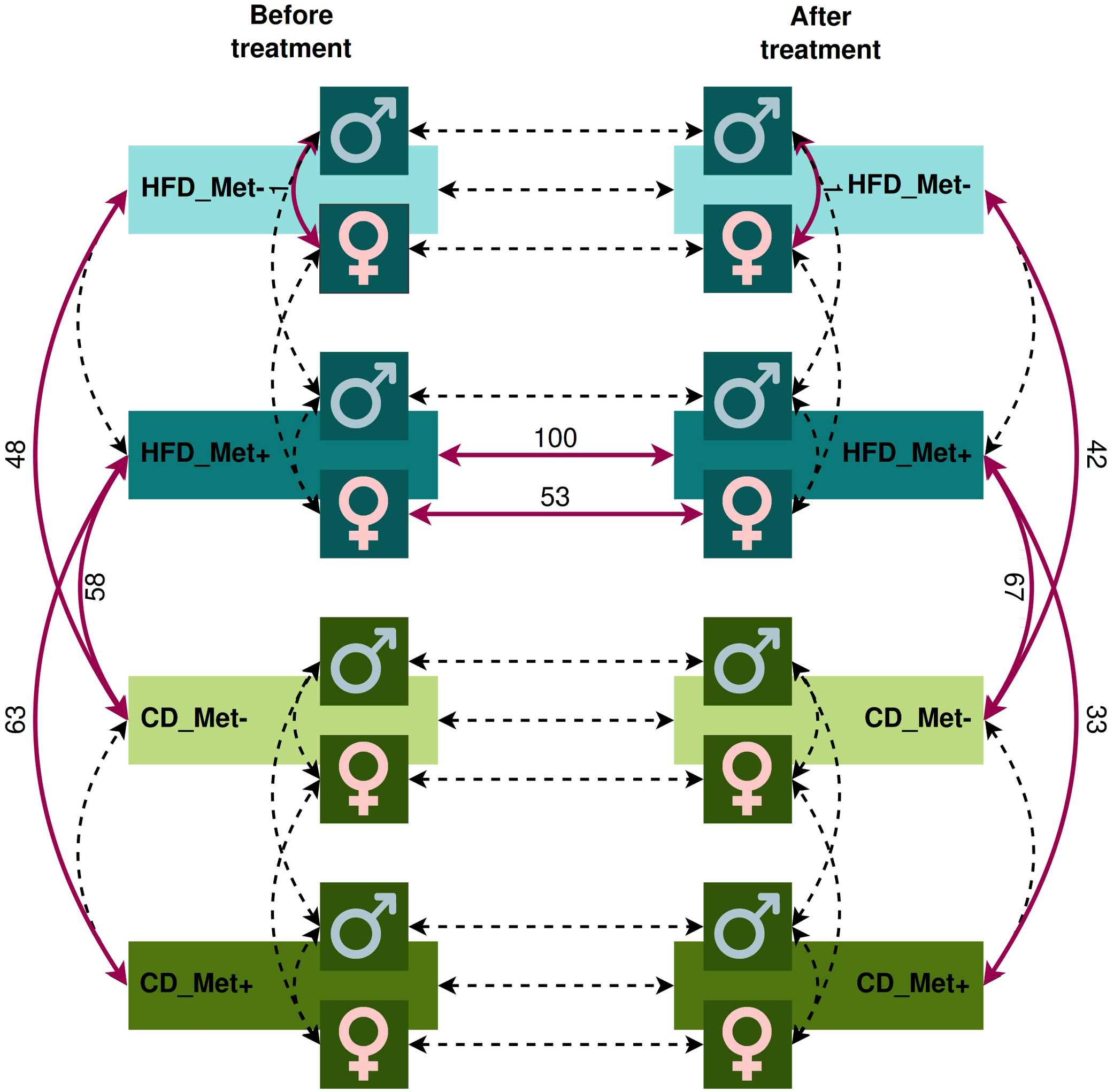
Contrasts in which microbiome compositions were compared. Dashed lines indicate the contrasts between which a comparison was performed. Red bold lines indicate the contrasts between which statistically significant differences in taxa abundance at species level were discovered with the numbers of the different species.

The analysis revealed significant differences between HFD_Met− and CD_Met− groups and between HFD_Met+ and CD_Met+ groups before the beginning of metformin treatment. We observed 63 and 48 differentially abundant species between HFD- and CD-fed groups with or without metformin treatment, respectively (Figure 6), indicating apparent differences between these experimental units at the time point when experimental units were allocated to receive metformin treatment. At the same time, no differentially abundant taxa were identified between CD-fed groups and between HFD-fed groups. When considering the effect of sex, only one species, *Bacteroides eggerthii* (LogFC = −4.05, FDR < 0.001), was detected to be differentially abundant between HFD-fed groups, indicating that some variability may exist between identically treated groups based on sex differences. No other substantial differences in microbiome composition between the same groups before the beginning of metformin treatment were found.

We did not find any differences between HFD_Met+ and HFD_Met− groups at the time point after the treatment (Figure 6). The same applies to the contrast between CD_Met+ and CD_Met− groups. When each of the sexes was compared, *Bacteroides eggerthii* remained significantly differentially abundant between HFD_Met− groups (LogFC = −3.64, FDR = 0.007).

We observed a strong effect of diet on microbiome composition both before and after metformin treatment. Before initiating metformin treatment, we observed 48 and 63 differentially abundant species in contrasts between HFD_Met− and CD_Met− groups and between HFD_Met+ and CD_Met+ groups, respectively. The same applied to identical contrasts at the end of the experiment; 42 differentially abundant species were identified between HFD_Met− and CD_Met− groups and 33 – between HFD_Met+ and CD_Met+ groups. Common to all of the contrasts mentioned above, HFD was associated with a lower abundance of *Bacteroidales* bacteria, *Prevotella sp.*, *Lactobacillus aviarius*, *Bacteroides helcogenes*, and *Bacteroides oleiciplenus*.

When comparing the differentially abundant taxa between HFD-fed and CD-fed groups representing the same metformin treatment status cross-sectionally, in all contrasts not influenced by metformin (HFD_Met− vs. CD_Met− before treatment, HFD_Met− vs. CD_Met− after treatment, and HFD_Met+ vs. CD_Met+ before treatment) in HFD groups we observed higher abundance of *Acetivibrio ethanolgignens* (LogFC values ranging from 3.07 to 3.84, FDR ≤ 0.008) and lower abundance of *Prevotella lascolai* (LogFC values ranging from −2.54 to −1.70, FDR ≤ 0.008), *Gabonia massiliensis* (LogFC values ranging from −3.09 to −1.77, FDR ≤ 0.02), *Culturomica massiliensis* (LogFC values ranging from −3.24 to −1.79, FDR ≤ 0.03), and several *Bacteroides* species (LogFC values ranging from −3.72 to −1.82, FDR ≤ 0.03).

The pairwise comparison between CD_Met+ groups before and after the treatment showed no differences in taxa abundance (Figure 6); the same was observed between CD_Met− groups.

Comparing HFD_Met− groups between the timepoints of metformin initiation and the end of the experiment, we observed no differentially abundant taxa. However, in HFD_Met+ groups, 100 species were altered (Figures 7a and 8), with most of the species abundancies being increased due to metformin treatment. The most pronounced changes were in the abundance of such *Bacteroidetes* genera as *Bacteroides*, *Parabacteroides*, *Prevotella*, *Paraprevotella*, *Porphyromonas; Firmicutes* genera *Bacillus*, *Butyrivibrio*, *Enterococcus*, *Lactobacillus*, *Lactococcus*, *Leuconostoc*, and *Streptococcus* as well as in *Enterorhabdus* representing *Actinobacteria*.

**Figure 7.**
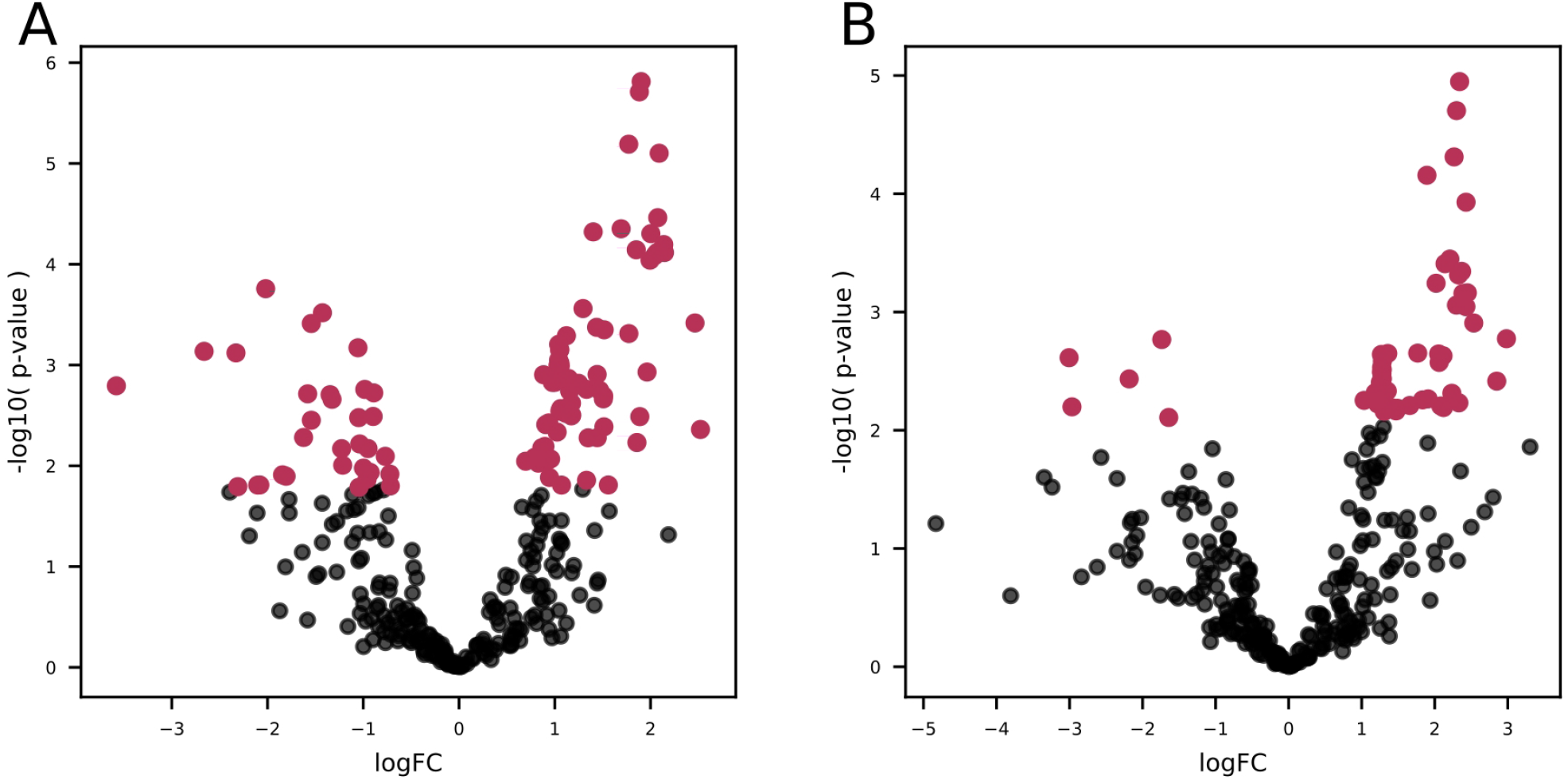
Volcano plots showing P-values of differentially abundant species in HFD_Met+ groups before and after metformin treatment. (**A**) P-values of differentially abundant species in HFD_Met+ groups before and after metformin treatment. (**B**) P-values of differentially abundant species in HFD_F_Met+ group before and after metformin treatment. Red dots represent differentially abundant species with P-values < 0.05.

**Figure 8.**
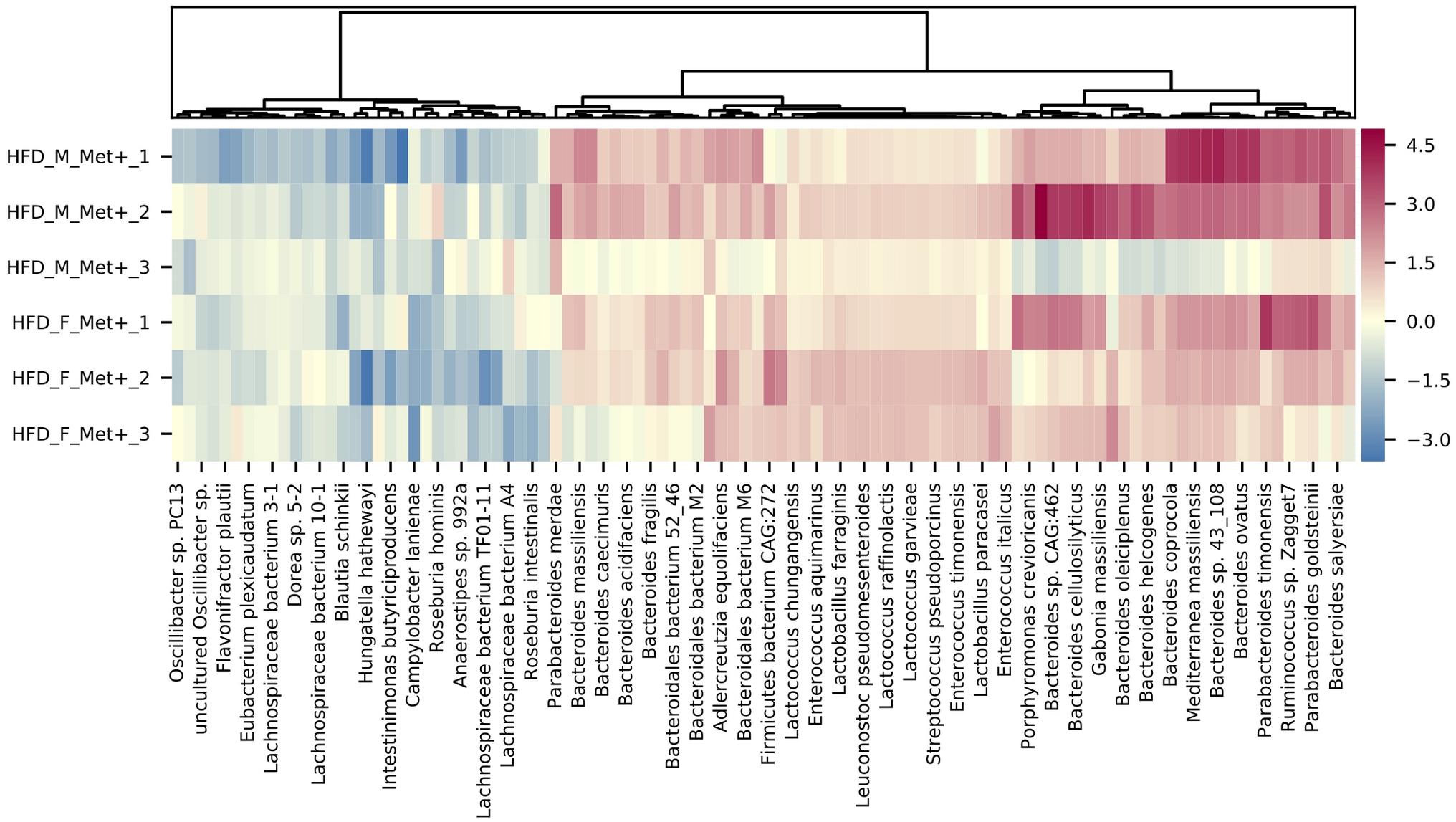
Heatmap showing differentially abundant species in HFD_Met+ groups before and after metformin treatment.

In general, the variability of the magnitude of the differences in the abundance of species between the samples of the studied contrast was more pronounced in species representing phylum *Bacteroidetes* (LogFC values ranging from 0.70 to 2.46). In contrast, members of the class *Bacilli – Lactobacillus*, *Lactococcus, Enterococcus, Leuconostoc, Bacillus*, and *Streptococcus* genera all representing *Firmicutes* were increased (LogFC values ranging from 0.79 to 1.17) in response to metformin in all of the samples relatively uniformly.

Analysis of taxa abundances in HFD_Met+ groups before and after metformin treatment in each of the sexes separately revealed sex-specific effects of metformin. In males, we did not identify significantly differentially abundant taxa while in females, the abundance of 53 species was significantly altered (Figures 7b and 9). A decrease in response to metformin was observed in bacteria of *Clostridia* class including *Faecalibacterium prausnitzii* (LogFC = −2.18, FDR = 0.03)*, Enterocloster clostridioformis* (LogFC = −1.74, FDR = 0.03) and *Anaerostipes sp. 992A* (LogFC = −1.64, FDR = 0.04) as well as *Desulfovibrio fairfieldensis* (LogFC = −2.97, FDR = 0.04) of *Deltaproteobacteria* class. We identified an increase in the differentially abundant taxa in response to metformin; a subtle increase in the abundance of species representing *Bacilli* was observed (LogFC up to 1.36), while the increase in *Bactoroidia* class representatives was particularly pronounced, for example, *B. ilei* (LogFC = 2.85), *B. vulgatus* (LogFC = 2.45), *B. pyogenes* (LogFC = 2.43). The list of differentially abundant species in females corresponds to the taxa identified in the analysis where samples from both sexes are taken together, although the extent of changes in abundances was greater in females.

**Figure 9.**
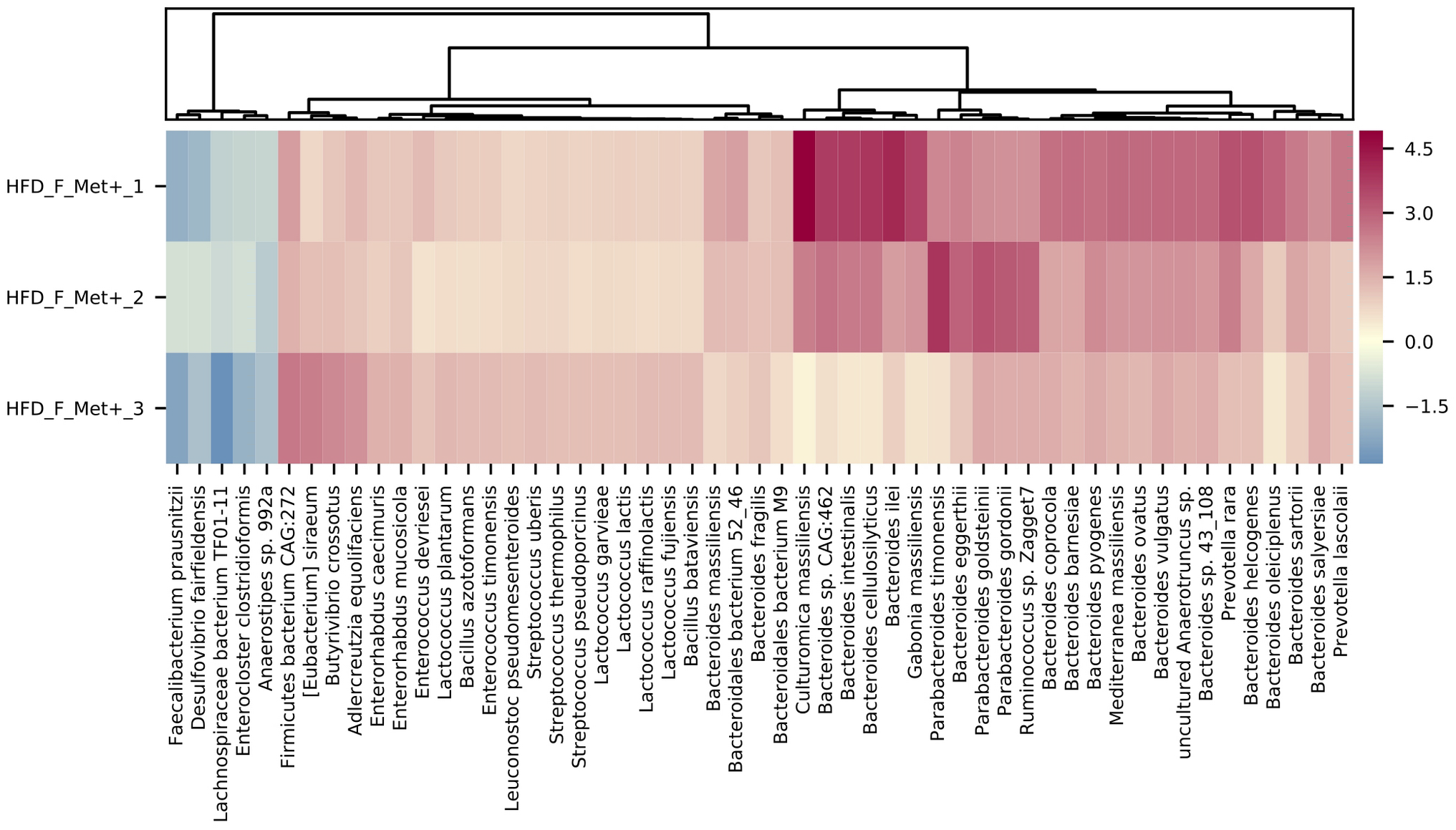
Heatmap showing differentially abundant species in HFD_F_Met+ group before and after metformin treatment.

To further analyze the effect of metformin treatment we compared microbial compositions between CD_Met− and HFD_Met+ groups. At the timepoint before the treatment we observed 58 differentially abundant taxa dominated by members of *Clostridia* and *Bacteroidetes* which were increased and decreased in HFD-fed animals, respectively. *Clostridiaceae* were represented by *Hungatella hathewayi* (LogFC = 6.65, FDR < 0.001) and *Acetivibrio ethanolgignens* (LogFC = 4.30, FDR < 0.001). Members of *Lachnospiraceae* including *Roseburia* and *Dorea* followed with LogFC from 0.94 to 3.35, FDR <0.05, as well as non-*Clostridia Mucispirillum schaedleri* (LogFC = 2.89, FDR = 0.02), and *Enterorhabdus mucosicola* (LogFC = 1.27, FDR = 0.02). *Bacteroidetes* were represented by various *Bacteroidales* with the most extreme changes identified in the abundance of *Bacteroidales bacterium M10* (LogFC = −10.33, FDR < 0.001), in addition *Parabacteroides timonensis* (LogFC = −2.04, FDR = 0.01), members of *Prevotella* (LogFC from −2.53 to −3.61, FDR < 0.001) and others were decreased.

The same contrast at the time point after the metformin treatment revealed 67 differentially abundant species. *Bacilli* and *Bacteroides* prevailed among the increased taxa, while *Clostridia* and different *Bacteroidetes* represented decreased taxa. *Bacilli* were represented by *Staphylococcus xylosus* (LogFC = 6.40, FDR = 0.02) and members of genera *Enterococcus*, *Streptococcus*, *Leuconostoc*, and *Lactococcus* (LogFC from 1.08 to 1.58, FDR > 0.05). LogFC of *Actinobacteria Enterorhabdus mucosicola* was 1.91, FDR = 0.004). Among increased *Bacteroides* were such species as *B. salyersiae*, *B. pyogenes*, *B. vulgatus*, *B. ovatus*, *B. coprocola*, *B. eggerthii*, *B. clarus*, *B. congonensis*, *B. caccae*, and *B. acidifaciens* (LogFC from 1.12 to 2.40, FDR < 0.05). Decreased *Bacteroidetes* included *Bacteroidales bacterium M10* (LogFC = −9.68, FDR < 0.001), other *Bacteroidales*, members of *Prevotella*, *Porphyromonas*, and *Bacteroides* species *B. ilei, B. helcogenes, B. gallinarum, B oleiciplenus, B. cellulosilyticus, B. togonis, B. intestinalis*, and *B. coprophilus* (LogFC from −1.19 to −5.62, FDR < 0.05). *Clostridia* were represented by *Dorea formicigenerans* (LogFC = −3.38, FDR = 0.04) and *[Eubacterium] infirmum* (LogFC = −4.65, FDR = 0.009).

The 58 % and 76 % of differentially abundant species were shared between contrasts HFD_Met− vs. CD_Met− and HFD_Met+ vs. CD_Met+, respectively. When the same contrasts were analyzed at the time point after the treatment, 83 % and 100 % were shared.

## Discussion

An effect of metformin on gut microbiome composition and function has been reported in previous studies both in humans and animal models and even *in vitro*. However, most of the studies used 16S rRNA gene sequencing and were limited to evaluating metformin effects on only one of the mice’s sexes. Our study’s strength is the inclusion of both sexes and the use of randomized block design. Our main aim was to reliably evaluate metformin’s effect on the microbiome rather than reach the maximal effect of metformin response with elevated doses. Therefore, we chose to use a smaller metformin dose (50 mg/kg body mass/day) compared to other studies (200-300 mg/kg body mass/day) to match the concentrations applied to humans. Besides, we fed the mice representing control groups in our study with a well-matched diet and not the regular chow, which is not standardized and has beneficial effects on the gut microbiome *per se* due to its short-chain fatty acid-increasing nutritional composition (Dalby et al., 2017). The use of regular chow is common in previous studies assessing the effect of metformin on the microbiome, which can substantially alter the results.

The results of our study are in line with previous research in this area. However, we found substantial sex-related differences in response to metformin treatment. In contrast to another study involving both sexes of mice where serum glucose level decreased after metformin treatment, especially in female mice, we observed an increase in plasma glucose level in females although male mice responded with a decrease in plasma glucose level in both HFD- and CD-fed groups (H. Lee, 2014). Also, insulin resistance indicated by the HOMA-IR index declined after the metformin intervention in HFD_M_Met+ group but, inconsistent with previously reported data, insulin resistance was augmented in HFD_F_Met+ group. The same pattern was manifested in CD_Met+ groups, thereby strengthening the discrepancies in metformin effects between sexes. These differences might be explained by the fact that we used a substantially lower dose of metformin (50 mg/kg body mass/day), which is considered the maximum dose that the body can utilize efficiently (He & Wondisford, 2015).

Intrinsic microbial diversity was higher in HFD-fed groups than CD-fed groups and decreased in response to metformin treatment in HFD-fed groups of both sexes but not in CD-fed groups in line with previous research (H. Lee, 2014). Beta diversity analysis revealed apparent clustering depending on diet and sex. The inclination toward *Lactobacillus* and *Proteobacteria* for HFD-fed female and male mice, respectively, at the time point before the treatment, coincides with the significant differences in glucose level and HOMA-IR between sexes at the same time point. This finding suggests that the abundance of *Proteobacteria* is associated with a higher level of insulin resistance, while *Lactobacillus* is negatively correlated with HOMA-IR.

Paired tests are more powerful in detecting microbial composition changes, explaining the fact that we observed 100 differentially abundant species comparing the samples before and after metformin treatment in HFD-fed groups. In contrast, we did not find significantly different taxa in the cross-sectional analysis comparing metformin-treated vs. untreated groups of HFD-fed mice at the time point after metformin treatment.

We observed that metformin influences the abundance of two *Firmicutes* classes in opposite directions, *Clostridia* being decreased or *Bacilli* being increased after metformin treatment. The abundance of *Bacteroidetes* and *Actinobacteria* was elevated, while all the members of *Proteobacteria* subsided in response to metformin.

We have observed a uniform effect of metformin treatment on bacteria, depending on their oxygen tolerance status and Gram staining. Metformin increased the abundance of facultatively anaerobic, aerotolerant, and obligate aerobic Gram-positive bacteria and obligate anaerobic Gram-negative bacteria. In turn, obligate anaerobic Gram-positive and microaerophilic and aerotolerant Gram-negative bacteria were decreased. As an exception, two genera representing *Eggerthellaceae* – *Enterorhabdus* and *Adlercreutzia* were increased despite being anaerobic and Gram-positive, which could be explained by their affiliation to another phylum, *Actinobacteria*.

We observed an increase in *Adlercreutzia equolifaciens* in both sexes, similarly to previously identified significant elevation of *Adlercreutzia* species in the fecal samples after metformin treatment. This increase has been explained by *Adlercreutzia*-mediated soybean isoflavonoid conversion to equol, which regulates glucose uptake in adipocytes by modulating known insulin-stimulation pathways (Napolitano et al., 2014).

Analysis of differences in microbiome composition between male and female mice showed divergence in taxonomic abundances changes. Similar to what has been reported previously, we observed a more pronounced increase in *Bacteroides* in female mice than male mice in HFD_Met+ groups (H. Lee, 2014). Genera representing *Firmicutes* were more consistent between both sexes, and in subjects, their abundance increased in response to metformin.

We compared CD_Met− and HFD_Met+ groups to investigate whether metformin shifts microbial composition in the direction of healthy, undisturbed microbiome expected to be observed in CD-fed groups not receiving metformin treatment. Metformin increased the abundance of *Bacilli* and some *Bacteroidetes*, in addition to a decrease of *Clostridia*. This finding supports previous reports that metformin shifts the dysbiotic microbiome associated with metabolic disease phenotypes to a microbiome corresponding to a more healthy state by increasing the abundance of members of *Bacteroidetes* (X. Li et al., 2017).

### Species with decreased abundance in response to metformin treatment

Species with the most substantially decreased abundance in response to metformin treatment *Hungatella hathewayi* (LogFC = −3.58, FDR = 0.01) has been previously described as a strictly anoxic, Gram-positive, spore-forming, rod-shaped bacterium isolated from human feces (Steer et al., 2001). The bacterium used carbohydrates as fermentable substrates, producing acetate, ethanol, carbon dioxide, and hydrogen as the major glucose metabolism products.

Members of *Proteobacteria* – sulfate-reducing bacteria *Desulfovibrio fairfieldensis*(LogFC = −2.66, FDR = 0.008)*, Desulfomicrobium orale* (LogFC = −2.08, FDR < 0.05) and *Mailhella massiliensis* (LogFC = −1.81, FDR = 0.04) and *Campylobacter lanienae* which has been isolated from fecal samples of asymptomatic individuals (LogFC = −2.10, FDR < 0.05) (Logan et al., 2000) were other species highly affected by metformin. *S*ulfate-reducing bacteria have been associated with inflammatory bowel diseases (IBD) (Loubinoux et al., 2002). It has been shown that metformin attenuates IBD severity and reduces inflammation (S. Y. Lee et al., 2015) suggesting a possible role of the microbiome in metformin effects on IBD.

Our study revealed a consistent effect of metformin on the abundancies of species representing *Clostridia*, more specifically members of *Lachnospiraceae* and *Ruminococcaceae*, as all of the identified differentially abundant taxa from these dominant butyrate-producing families (except for *Butyrivibrio* and *Rumonicoccus sp. Zagget7*) were decreased after the treatment when analyzing the HFD_Met+ group longitudinally. Butyrate-producing bacteria *Roseburia hominis*, *Roseburia intestinalis, Faecalibacterium prausnitzii* – dominant butyrate-producing *Firmicutes* found in the human intestine (La Rosa et al., 2019), *Intestinimonas butyriciproducens* and *Eubacterium plexicaudatum* first found in mouse intestine (Kläring et al., 2013; Wilkins et al., 1974) and *Anaerotruncus spp.* are among these taxa. Butyrate has been demonstrated to positively impact gastrointestinal tract homeostasis, as it promotes the growth of intestinal epithelial cells, increases the expression of tight junction proteins, and acts as an anti-inflammatory agent (Kant et al., 2016). Previous studies have reported that obesity and T2D are associated with gut microbiome dysbiosis, a reduction in butyrate-producing bacteria and an increase in opportunistic pathogens (Forslund et al., 2015; McCreight et al., 2016). A recent randomized pilot study in which the impact of probiotic supplement on metformin’s effect on glycaemia in prediabetic subjects was assessed, has identified an increase in the abundance of *Anaerotruncus colihominis* (Cluster IV) only in participants receiving both metformin and the probiotic but not in participants taking either metformin or probiotic alone (Palacios et al., 2020). Our results show that metformin alone can impact the abundance of different *Anaerotruncus* species in opposite directions; *Anaerotruncus sp.* G3(2012) was decreased, while the abundance of uncultured *Anaerotruncus sp.* was augmented. In contrast to our findings, the potential role of *Blautia* species in the maintenance of a metabolically healthy phenotype and the management of obesity, insulin resistance, and T2D has been proposed (Benítez-Páez et al., 2020). Another member of *Lachnospiraceae – Dorea* is shown to be increased in T2D individuals and negatively correlated with the abundance of butyrate-producing bacteria (Q. Li et al., 2020). According to our data, this observation propounds the idea that *Dorea* is negatively correlated with butyrate producers representing *Bacteroidia* but not *Clostridia*. Gene-targeted approaches to investigate the butyrate-producing bacterial communities of the human gut microbiome have suggested that butyrate-producing colon bacteria form a functional group rather than a monophyletic group (Rivière et al., 2016). This finding suggests that the functional niche of *R. hominis*, *R. intestinalis*, *I. Butyriciproducens*, and *F. prausnitzii* as butyrate-producing bacteria is substituted by other taxa, possibly butyrate-producing species identified with an increased abundance in response to metformin treatment which was discovered in our study.

A recent study has discovered *Lachnoclostridium* sp. members as fecal bacterial markers for early detection of colorectal cancer by being significantly enriched in adenoma patients (Liang et al., 2019). Another species with decreased abundance in response to metformin is *Flavonifractor plautii*, a flavonoid-degrading bacteria that has been associated with colorectal cancer in Indian patients (Gupta et al., 2019). Flavonoids are substantial components of the human diet and have favorable effects on the prevention of T2D (Del Rio et al., 2013). Flavonoids could be affecting the composition of the gut microbiome by promoting beneficial bacteria and preventing the increase of potential pathogens (Braune & Blaut, 2016). The latest research has indicated that metformin has benefits in diabetes treatment and in lowering the risk of developing cancer, including colorectal cancer (Higurashi & Nakajima, 2018). Our findings suggest that chemoprevention effects of metformin could be mediated through the gut microbiome.

### Species with increased abundance in response to metformin treatment

This study revealed a uniform effect of metformin treatment on bacteria representing *Bacilli*. Genus *Lactobacillus* is widely represented among the increased taxa by *L. murinus*, *L. farraginis*, *L. paracasei*, *L. kefiranofaciens*. This finding is consistent with previous work showing that metformin increases the abundance of genus *Lactobacillus* in HFD-fed rats (Bauer et al., 2018). Probiotic supplementation with *Lactobacillus* improves glucose parameters in diabetic rats and prevents insulin resistance and hyperglycemia in HFD-fed mice (Bauer et al., 2018), which is in line with our observations. The abundance of another genus of lactic acid bacteria *Lactococcus*, including *L. raffinolactis*, *L. garvieae*, *L. fujiensis*, *L. chungangensis, L. lactis, L. plantarum* is increased in response to metformin treatment, which is in agreement with a recent study where the effect of metformin on short term HFD-induced obesity-related gut microbiome was evaluated (Ji et al., 2019).

Previous work has highlighted that metformin increases gut utilization of glucose and subsequent lactate production, which may be due to metformin’s action on the gut microbiome (McCreight et al., 2016). Local lactate concentrations might be shaping the gastrointestinal symptoms associated with metformin intolerance (McCreight et al., 2016).

Similarly to the identified differences in *Streptococcus* in response to gut microbiome modification by HFD and metformin (Ji et al., 2019), we observed a consistent increase in several species representing the genus in HFD_Met+ groups of both sexes. Heat-killed *S. thermophilus* has been shown to moderate insulin resistance and glucose intolerance and protect the intestinal barrier in the ZDF T2D rat model (Gao et al., 2019).

One of the gut microbiome’s main roles is to break down the dietary fiber and starch incompletely hydrolyzed by the host. Short-chain fatty acids (SCFA), including propionate and butyrate, are the main fermentation products of fiber digestion and can be used for lipid or glucose *de novo* synthesis. Changes in the SCFA profiles are associated with changes in the gut microbiome (McKnite et al., 2012).

In our study, taxa associated with SCFA production, such as *Bacteroides* and *Enterorhabdus*, which convert amino acid derivatives as the energy source (Clavel et al., 2014), are represented among the species with increased abundance after metformin treatment.

*Enterorhabdus* has been represented exclusively in lean mice when compared with db/db diabetic mice (Geurts et al., 2011). Moreover, Clavel et al. have identified that members of the *Coriobacteriaceae*, with the main species including *Enterorhabdus mucosicola*, *Enterorhabdus cecimuris*, in the cecum can be related to obesity resistance (Clavel et al., 2014). Our data support these reports as previous studies have associated body weight loss with metformin therapy (Yerevanian & Soukas, 2019).

The abundance of *Bacteroides* was significantly higher after the metformin treatment, with the most represented bacteria being *B. intestinalis*, *B. vulgatus*, and *B. acidifaciens*. However, the effect was heterogeneous, with an observed decrease of several *Bacteroides* species in some experimental units. The increased abundance of *Bacteroides* and *Parabacteroides* is in line with existing data (H. Lee et al., 2018).

It has been described that *B. fragilis* colonization aggravates metabolic disorders induced by HFD, whereas metformin inhibits the growth of *B. fragilis* through modification of folate and methionine metabolism (Sun et al., 2018). In contrast, our study has shown an increase in *B. fragilis* abundance in response to metformin treatment in HFD_Met+ groups of both sexes.

In a study on core gut bacteria of healthy mice, a high prevalence (99%) and a relatively high abundance of *Parabacteroides* was observed and suggested that *Parabacteroides* might be essential to host health (Wang et al., 2019). This fact corresponds to metformin’s reported health benefits regarding the microbiome composition shift (Wu et al., 2017). It has been hypothesized that the microbial growth-inhibitory effect of metformin is more pronounced on anaerobic organisms than aerobic organisms as anaerobic respiration produces less ATP than aerobic respiration (Prattichizzo et al., 2018).

Metformin-induced changes in the abundance of *Akkermansia muciniphila* and metabolic improvement due to these changes have previously been shown in several studies (H. Lee, 2014; Wu et al., 2017). Instead, our study reports no significant changes in *A. muciniphila* abundance in response to metformin in fecal metagenomic data.

Differences in experimental designs can explain discrepancies in results between our study and previously reported ones. Most of the studies have paid less attention to properly define the identified experimental unit in their experiments, which in microbiome studies most often should be a cage with animals, leading to pseudoreplication. This practice potentially evokes the exaggeration of the identified differences in data. Even though we performed sample size estimation before this complex experiment, this study’s results would benefit from analysis in larger groups as the variability of metabolic parameters and gut microbiome composition was noteworthy.

In conclusion, our study identified changes in microbial diversity and composition of the gut microbiome in response to metformin treatment in high fat diet-fed mice. Furthermore, sex-specific differences were discovered where male mice experience more pronounced changes in metabolic markers; however, inverse effects on metabolic markers were revealed in female mice together with more marked changes in gut microbiome composition. We suggest that both sexes of animals should be included in future studies focusing on metformin effects on the gut microbiome.

## Author Contributions

LS designed and performed the experiment, collected samples, prepared samples for sequencing, wrote the manuscript. IS performed data processing and analysis. MU and IK performed the experiment. ZK, RP and IE collected samples. JK supervised the study and revised the manuscript. All authors gave final approval and agree to be accountable for all aspects of the work.

